# System-wide quantitative proteomics of the metabolic syndrome in mice: genotypic and dietary effects

**DOI:** 10.1101/051508

**Authors:** Camille Terfve, Eduard Sabidó, Yibo Wu, Emanuel Gonçalves, Meena Choi, Stefania Vaga, Olga Vitek, Julio Saez-Rodriguez, Ruedi Aebersold

**Author notes:** Contributed equally to this work (Camille Terfve, Eduard Sabidó, Yibo Wu).

## Abstract

Advances in mass spectrometry have made the quantitative measurement of proteins across multiple samples a reality, allowing for the study of complex biological systems such as the metabolic syndrome. Although deregulation of lipid metabolism and increased hepatic storage of triacylglycerides are known to play a part in the onset of the metabolic syndrome, its molecular basis and dependency on dietary and genotypic factors are poorly characterized. Here, we used a rich experimental design with two different mouse strains, dietary and metabolic perturbations to generate a compendium of quantitative proteome data, using three mass spectrometry strategies. The data recapitulates known properties of the metabolic system and indicate differential molecular adaptation of the two mouse strains to perturbations, contributing to a better understanding of the metabolic syndrome. We show that high-quality, high-throughput proteomic datasets provide an unbiased broad overview of complex systems upon perturbation.

**HIGHLIGHTS:**

- Rich experimental design with two mouse strains, dietary and metabolic perturbations
- Three mass spectrometry proteomic data collection strategies in one data compendium
- SWATH-MS data with systematic perturbation provide a broad unbiased view of the system
- The data reveals differential adaptation of molecular systems in two mouse strains

## INTRODUCTION

Most biological processes are complex systems consisting of multiple molecules such as DNA, transcripts, metabolites and proteins, and their interplay and dynamic interactions eventually determine the observed phenotypes [1]. A common approach to study such systems is to quantify the elements of the system under a number of perturbed conditions using high-throughput methods and to integrate the data into dynamic network models, or to generate hypotheses that are then followed up using traditional molecular biology methods [2]. To date most of these measurements have been carried out at the transcript level due to the maturity of the technology. However, because proteins are closer to biological function than transcripts and their levels not necessarily strongly correlated with transcript levels [3], it can be expected that complementary information on the system can be gained by additional proteome measurements.

Broadly, two proteomics approaches have been used to measure proteomes: discovery and targeted proteomics. In discovery or shotgun proteomics, protein mixtures are enzymatically digested and the generated peptides are then separated by liquid chromatography and subsequently analyzed by tandem mass spectrometry. Peptide sequences are deduced from experimental spectra, and protein quantities are inferred from the signal intensity of the corresponding peptide ion, using different heuristics. Both proteome coverage and accuracy of discovery (equivalently, shotgun) proteomics approaches have improved tremendously over the last few years, and it is still the most popular mass spectrometric technology to study proteomes [4, 5]. However, due to the underlying algorithms used to select peptide ions for fragmentation, discovery shotgun proteomics suffers from under-sampling, and the overlap between proteins identified across samples is insufficient, particularly, if highly complex samples are being analyzed. Therefore, the number of conditions that can be meaningfully compared with this approach is limited.

In contrast to discovery proteomics, targeted proteomics approaches such as selected reaction monitoring (SRM) quantify a reduced set of pre-determined proteins of interest across sample sets. In the SRM approach, peptide precursor ions corresponding to the targeted protein are isolated and fragmented, and then a pre-defined fragment ion is further selected and its signal recorded over time. Each pair of precursor and fragment ion is called a transition and multiple transitions are coordinately measured and used to identify and quantify the selected peptide in a complex sample. Indeed, when used to quantify a selected subset of proteins, SRM results in near complete data matrices (proteins quantified across conditions), higher dynamic range, and higher quantification precision when compared to classic shotgun proteomics [6]. However, because the number of proteins quantified per analysis is limited and needs to be designated prior to the measurement, the generated results strongly depend on the *a priori* selection of target proteins by expert knowledge. This constraint contrasts with the capabilities of screening approaches such as discovery proteomics that are not limited by the selection of a subset of proteins of interest, and thus, permits protein quantification in complex samples without any previous requirement of expert knowledge [4, 5, 7].

Recently we have developed SWATH-MS, a mass spectrometric method that combines some of the strengths of each method by combining the high-throughput capabilities of discovery approaches with the consistent quantification exhibited by SRM [8]. The method is based on the acquisition of fragment ion spectra of all precursor ions within a narrow mass-to-charge window on a high-resolution MS instrument, where the entire mass-to-charge space is repeatedly scanned across the chromatographic retention time space. The complex fragment ion spectra and retention time information can then be queried for the presence and quantity of peptides of interest using *a priori* information such as fragmentation pattern and retention time. Our group has previously shown the quantitative accuracy achieved by this method to be comparable to that of SRM, with sensitivity at an intermediate level between SRM and shotgun approaches, but with proteome coverage capabilities at least equal to shotgun and the possibility to virtually endlessly re-mine the data for new peptides after mass spectrometric acquisition [8].

In this paper we applied the three aforementioned proteomic strategies, i.e. discovery or shotgun proteomics, selected reaction monitoring (SRM), and SWATH-MS to the analysis of a systematically perturbed complex biological system in the context of the metabolic disease syndrome. The metabolic syndrome encompasses a series of disorders, including obesity, insulin resistance and hepatic steatosis, which are associated with predisposition to diabetes, cardiovascular disease and cancer. The prevalence of this disease has increased to unprecedented levels in the last decades, fostering much interest in studying the relationship between these pathologies and the molecular mechanisms underlying its etiology [9, 10, 11]. Deregulation of lipid metabolism and increased storage of triacylglycerides in the liver have been proposed as the main processes involved in the development of the metabolic syndrome and its link to the onset of insulin resistance [9, 12]. Indeed, a series of studies have investigated the phenotypic, transcriptional and/or proteomic response of different mouse strains to long-term high-fat diet [9, 10, 11, 13], but the connection between these metabolic alterations is not yet fully understood, and neither is the influence of genetic and environmental factors on the establishment of the metabolic syndrome [9, 13]. We show that targeted high-throughput approaches such as SWATH-MS can quantify a larger number of proteins than SRM, and that overall the quantified proteins have similar variability to that of SRM. Moreover, we report a set of biological findings that describe differential molecular behaviors between the mouse strains, and contribute to a better understanding of the mechanisms of the metabolic syndrome and its link with the genetic and environmental factors.

## RESULTS

### A mass spectrometry proteome compendium

Liver samples from C57BL/6J and 129Sv mice (genetic background) fed with a high-fat rodent diet for 0, 6 and 12 weeks (time) were collected in biological triplicate either after *ad libitum* feeding or after overnight fasting (treatment), making a total of 36 samples for 12 different experimental conditions (**Figure 1**). C57BL/6J and 129Sv mouse strains were chosen for this study because of their distinctive phenotypic differences in response to a sustained high-fat diet, with C57BL/6J mice showing higher lipid accumulation in the liver, higher weight increase, a pronounced hyperglycemia and higher hyperinsulinemia when compared with the 129Sv strain [9, 12, 13].

**Figure 1.**
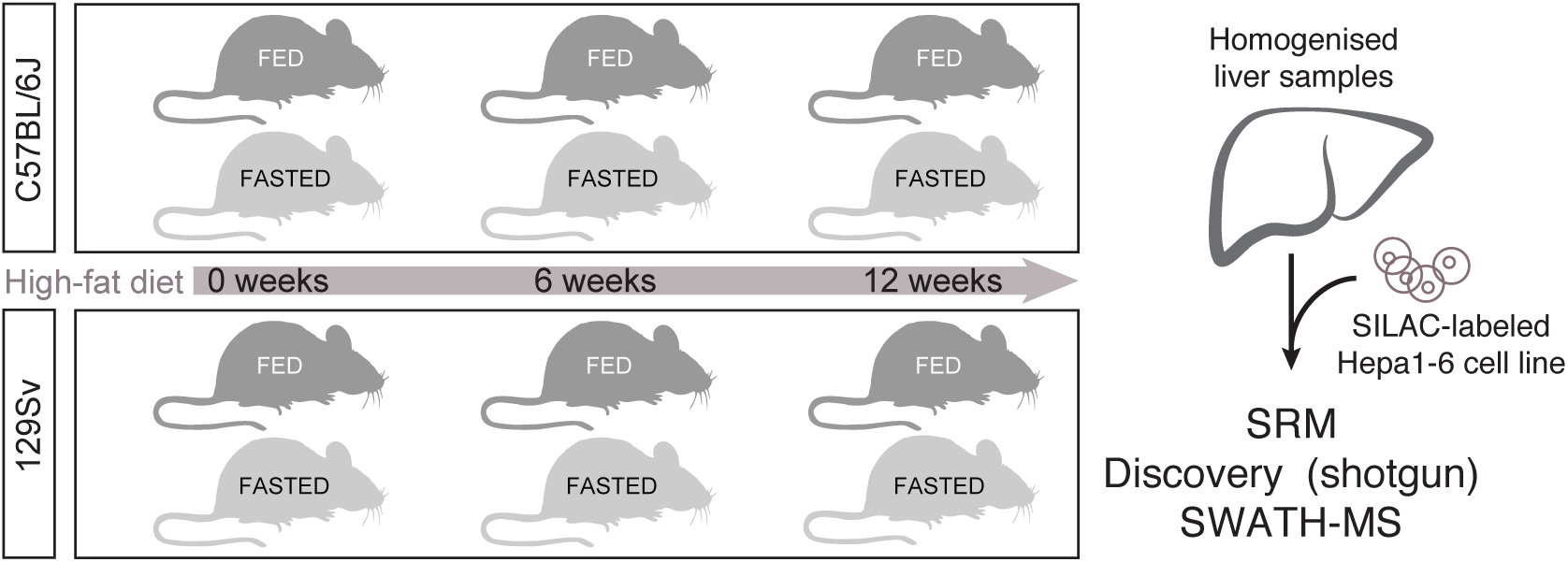
Experimental design. Two mouse strains that show phenotypically different responses to sustained high-fat diet were fed with a high-fat diet for 0, 6 or 12 weeks, with or without overnight fasting prior sacrifice. Mouse livers were extracted, homogenised and digested with trypsin for mass spectrometric analysis with three different mass-spectrometric approaches: discovery proteomics, selected reaction monitoring (SRM), and SWATH-MS. An isotopically-labelled hepatocyte HepG2 cell-line was used as a spiked-in reference for quantitation.

Collected liver samples were homogenized and digested with trypsin, and the resulting peptide mixes were subjected to LC-MS/MS analysis with different approaches: i) peptide fractionation followed by discovery shotgun proteomics analysis, hereafter referred to as *discovery dataset* ii) peptide fractionation followed by targeted proteomics analysis via selected reaction monitoring (SRM) of proteins associated with the insulin-signaling pathway and central metabolism, hereafter referred to as *the SRM dataset* [12]; and iii) SWATH-MS of unfractionated protein homogenates. These analyses allowed the consistent quantitation across all or most experimental conditions of 1235, 122 and 2203 proteins for the *discovery, SRM*, and *SWATH-MS* datasets, respectively (**Supplementary Table ST1**). The overlap of proteins was of 1114 between SWATH and shotgun, 78 between SWATH-MS and SRM, and 60 between SRM and shotgun (**Supplementary Figure S1A**).

We initially assessed the information content in the three acquired datasets by testing the proteins for differential abundance across all pairwise comparisons between time points within each mouse strain. Specifically, we compared i) B6T1FED–B6T0FED, ii) B6T2FED–B6T0FED, iii) B6T2FED–B6T1FED, iv) S9T1FED–S9T0FED, v) S9T2FED–S9T0FED, and vi) S9T2FED–S9T1FED, where B6 and S9 represent C57BL/6J and 129Sv mice, respectively, fed ad libitum (FED), after 0, 6 and 12 weeks (T0, T1 and T2) of high-fat diet. **Figure 2** **and Supplementary Figures S1B and S1C** show the results for all the proteins in the datasets, and **Supplementary Figure S2** shows the results for the subset of overlapping proteins acquired in all the datasets.

**Figure 2.**
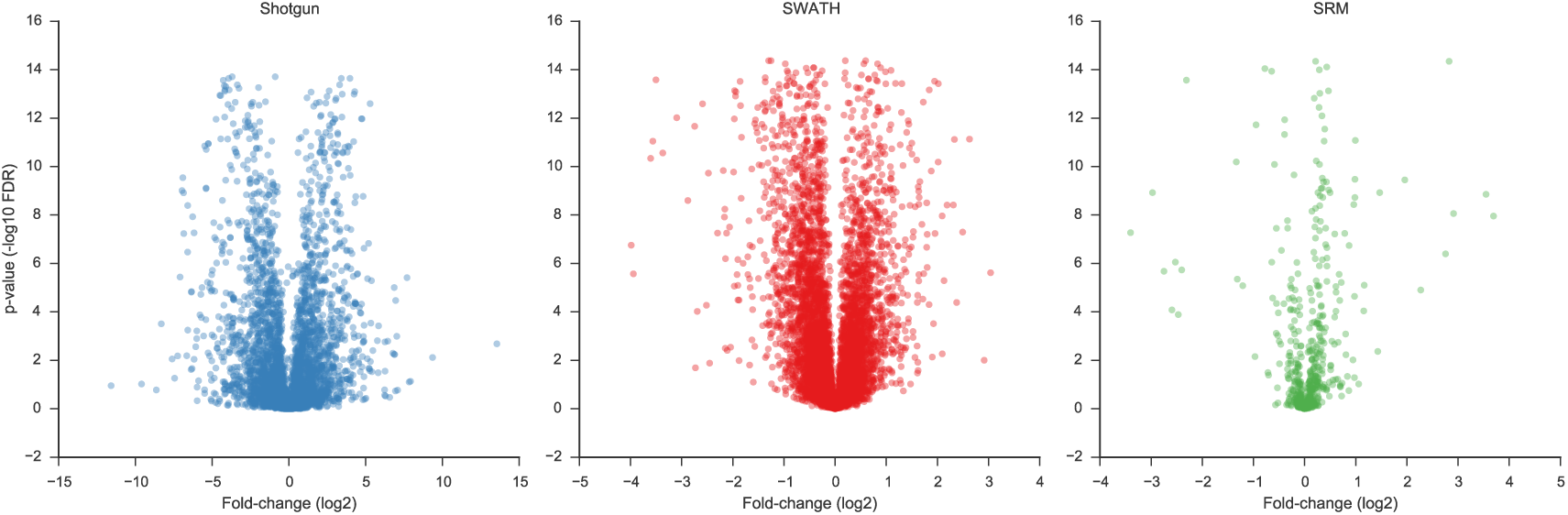
Distributions of the data in each data set. Volcano plots representing the-log10 of the adjusted p-value versus the log2-fold-change protein estimates for the discovery shotgun (a), SWATH-MS (b) and SRM (c) datasets. Each plot shows the overlayed values for the following tested comparison between conditions: i) B6T1FED–B6T0FED, ii) B6T2FED–B6T0FED, iii) B6T2FED–B6T1FED, iv) S9T1FED–S9T0FED, v) S9T2FED–S9T0FED, and vi) S9T2FED–S9T1FED). B6 represents C57BL/6J mice, and S9 represents 129Sv mice. T0, T1 and T2 correspond to 0, 6 and 12 weeks of high-fat diet. FED stands for mice that were fed *ad libitum* (treatment).

The patterns of the volcano plots correspond to our prior expectations from these technologies. In discovery shotgun, the distribution of log-fold changes of each condition to the reference state is wider than in the other two technologies, and there are more extreme values. However, fewer of the high-magnitude log-fold changes (61.0 % of those with fold-change larger than 2 have an adjusted p-value < 0.01) are statistically significant, in contrast to 97.2 % for SWATH or 100.0 % for SRM (see **Figure 2**). This is the effect of a relatively large technical variation associated with the shotgun workflow. In SRM, log-fold changes of large magnitude have a much stronger statistical evidence of differential abundance (i.e., small p-values). This is due to both relatively low technical variation, and to a careful choice of protein targets for SRM quantification. The SWATH-MS dataset can be thought of as an intermediate case. The volcano plot is broader in the bottom than SRM (indicating that some large magnitude log-fold changes are not statistically significant), however it has more proteins with strong evidence for differential abundance than the discovery shotgun dataset (indicating less technical noise). Overall, the log-fold changes distribution of SWATH-MS resembles SRM more than Shotgun (**Supplementary Figures S2**), suggesting that SWATH-MS acquisition can provide data with less technical variability and with data quality comparable to that generated by traditionally targeted approaches such as SRM. This is in agreement with data obtained in less complex proteome samples where comparable quantitative accuracy was obtained with SRM and SWATH-MS [8].

We further evaluated the Pearson correlations of protein log-fold changes between pairs of acquisitions, among the proteins quantified in both pairs (**Figure 3** **and Supplementary Figure S3**). The correlations between SWATH-MS and SRM were higher than between Shotgun and SRM. Moreover, a principle component analysis (PCA) was performed using the protein abundances per condition as estimated by MSstats for the three datasets. The results showed the ability of this analysis to distinguish among the different factors tested in the experiment (**Figure 4**). In all the cases, visualization of each experimental tested condition in the space of the first and second principal components produced clustering of strains (genetic background), but only the SRM and SWATH-MS datasets could separate the time factor using the combination of the first and third principal component. While the SWATH-MS measurements provided a more clear separation of the time factor than the SRM data-set, in SRM the first 3 principle components explained a larger proportion of the total between-sample variation across the samples. SWATH-MS and Shotgun had more unexplained variation, reflecting the challenges of finding signals in a larger protein space.

**Figure 3.**
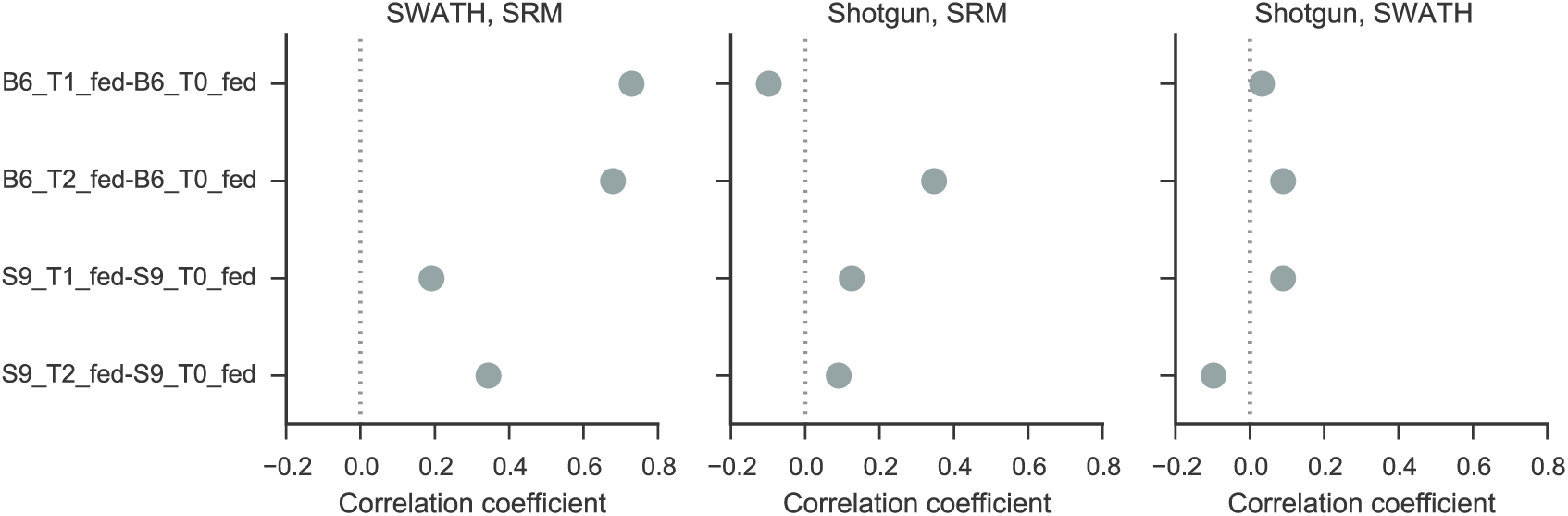
Analysis of correlation patterns between the different datasets. Pearson correlation coefficients between the obtained protein fold-changes in the discovery shotgun, SRM and SWATH-MS datasets for different comparisons (T0=0 weeks; T1=6 weeks; and T2=12 weeks of sustained high-fat diet).

**Figure 4.**
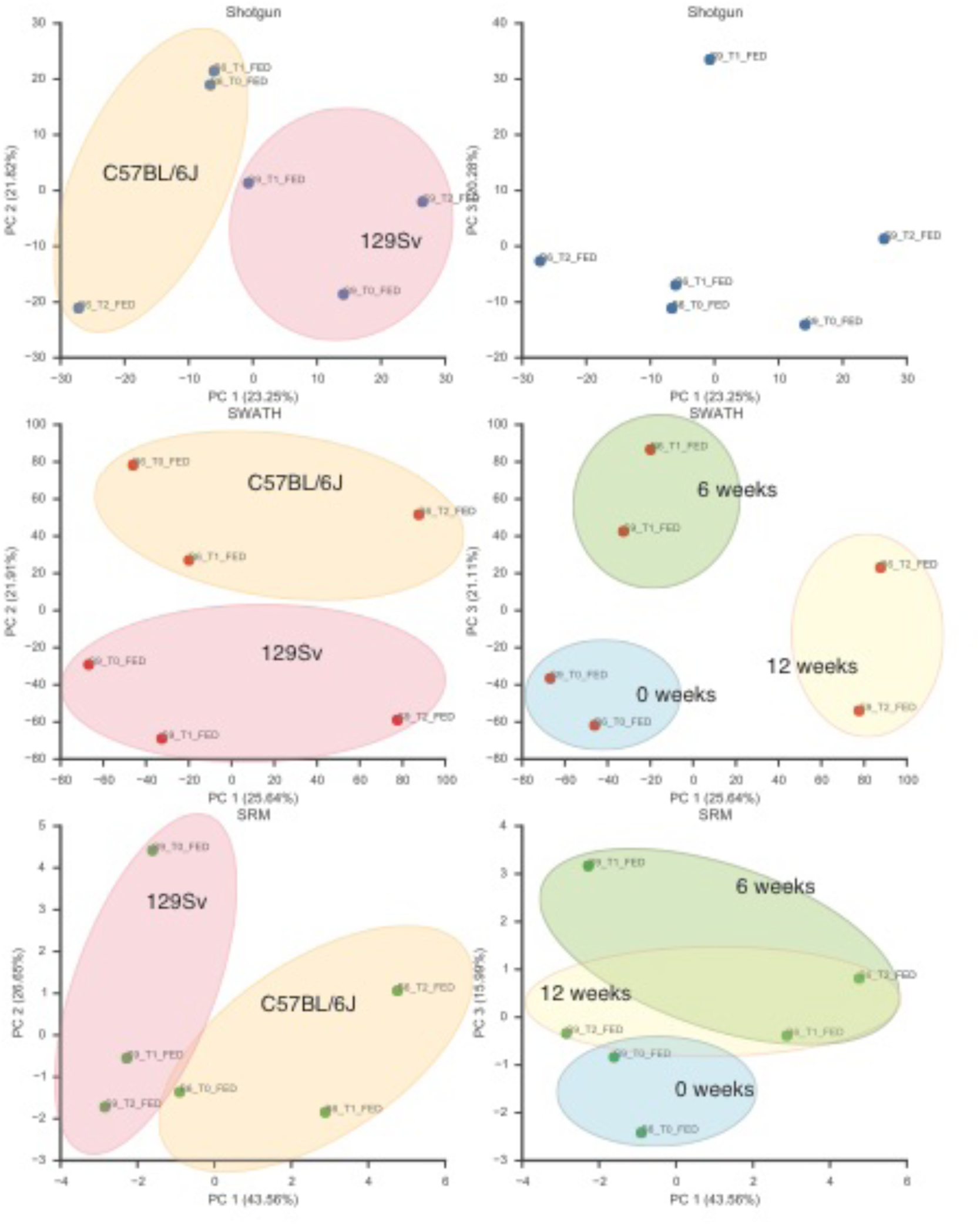
Principal component analysis of the individual datasets. Principal component analysis using protein abundance estimates per condition obtained by MSstats for the discovery shotgun, SRM and SWATH-MS. The % in each axis denotes the percentage of the sum of the protein variances across the samples, which is explained by the corresponding principal component.

### Time effects of a sustained high-fat diet on different genetic backgrounds

Based on the capability to quantify thousands of proteins with similar performance as SRM, the SWATH-MS dataset was selected for further downstream analysis to study the effects of the *genetic background* in the response to a sustained high-fat diet of 0, 6 and 12 weeks *(time)* in the studied mouse strains (C57BL/6J and 129Sv). Compared with previous targeted studies [12], with more than 2000 proteins consistently quantified across the conditions the SWATH-MS dataset allowed us to obtain a broader view of the cell state after long periods of high-fat diet, and it enabled to explore the remodeling of biological process beyond the insulin-signaling pathway and the central metabolism (**Supplementary Figure S4**).

Initially, the acquired dataset was used to evaluate the strain-specific protein responses after 6 and 12 weeks of high-fat diet *ad libitum* by comparing the fold-changes of a given protein between two time points in C57BL/6J with the protein fold-change between the same time points in 129Sv mice. This generated three main comparisons to be assessed: i) (B6T2FED–B6T1FED)–(S9T2FED–S9T1FED); ii) (B6T2FED–B6T0FED)–(S9T2FED–S9T0FED); and iii) (B6T1FED–B6T0FED)–(S9T1FED–S9T0FED); with B6 representing in C57BL/6J mice, S9 representing 129Sv mice, and T0, T1, and T2 corresponding to 0, 6 and 12 weeks of high-fat diet. In all cases the mice were fed *ad libitum* (treatment *FED*). Out of these analyses, we identified 964 proteins exhibiting a statistically significantly change in protein abundance in response to a sustained high-fat diet in at least one time point between mouse strains (adjusted p-value ≤ 0.01; Supplementary Table ST2). Among proteins with strain-specific response to high-fat diet, we identified several proteins from glycolysis and gluconeogenesis (HXK1, HXK4, F16P1, ALDOB, ALDOC, G3P, PGAM1, ENOA, KPYR, PCKGC, PYC), fatty acid metabolism (FAS), and beta-oxidation (ACOX1, ACADM, ECHP, DHB4) in agreement with previous results obtained by targeted quantitation of a selected subset of proteins by SRM [12] (**Figure 5**). For instance, we reproduced the time profiles for fatty acid synthase (FAS), a central enzyme in the fatty acid biosynthesis metabolism, and corroborated that about half of the proteins exhibiting different time responses to a sustained high-fat diet already diverged after 6 weeks (483 proteins; adjusted p-value ≤ 0.01), as it has also been previously described by protein quantitation with SRM [12]. Together, these results show that targeted high-throughput approaches such as SWATH-MS are able not only to distinguish between biologically relevant factors in complex experimental designs, but also to recapitulate the findings obtained with gold-standard methods in quantitative proteomics, i.e. SRM, without the need of any previous expert knowledge for target protein selection before data acquisition.

**Figure 5.**
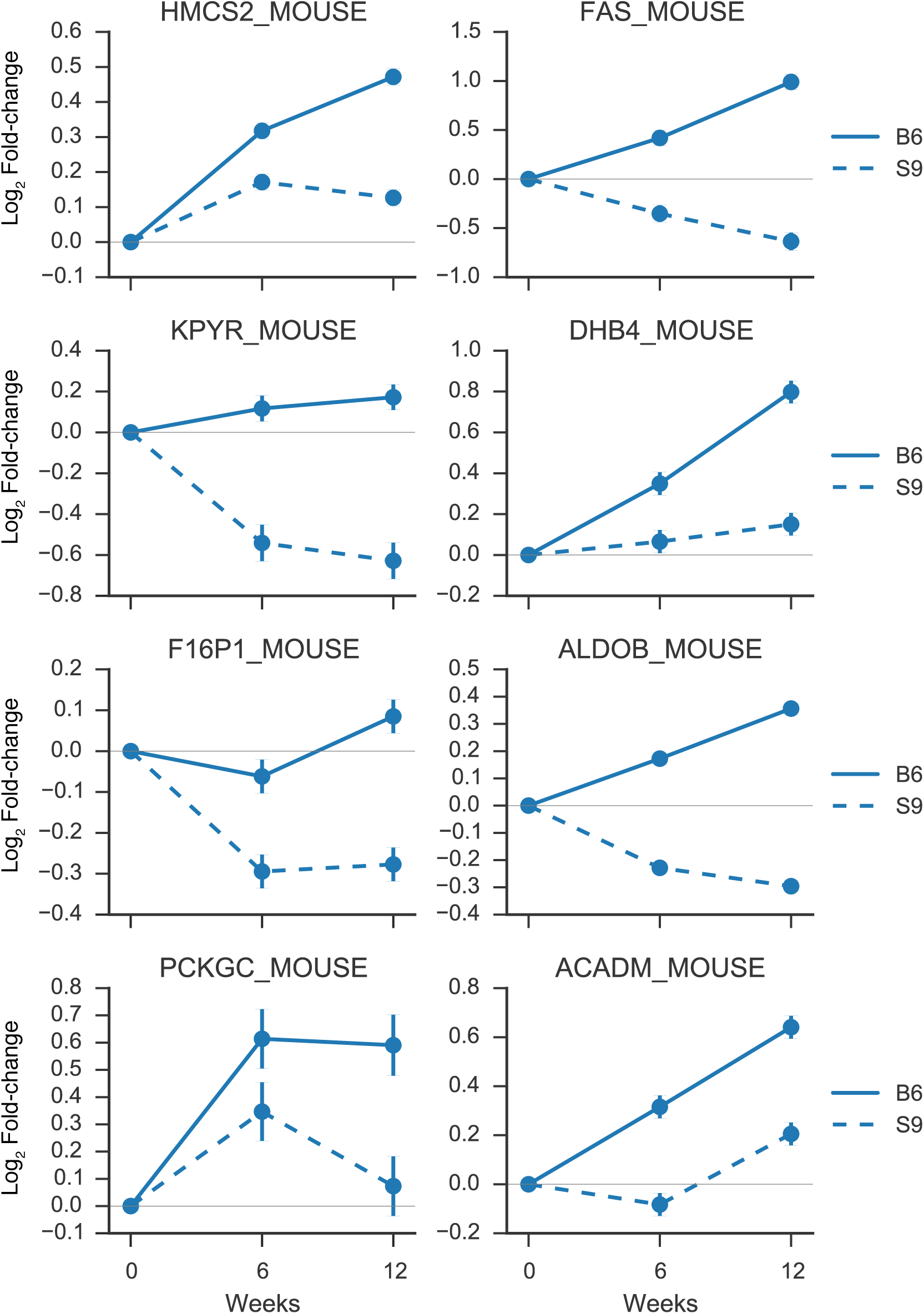
Protein profiles in response to several weeks of high-fat diet. Fold-changes (log-scale) corresponding to selected proteins involved in the main metabolism after 0, 6 and 12 weeks of sustained high-fat diet for C57BL/6J (B6) and 129Sv (S9) mice in the SWATH-MS quantitative dataset i.e. the three main comparisons that were assessed were: i) (B6T2FED–B6T1FED)–(S9T2FED–S9T1FED); ii) (B6T2FED–B6T0FED)–(S9T2FED–S9T0FED); and iii) (B6T1FED–B6T0FED)–(S9T1FED–S9T0FED); with B6 representing in C57BL/6J mice, S9 representing 129Sv mice, and T0, T1, and T2 corresponding to 0, 6 and 12 weeks of high-fat diet.

Next, we annotated the proteins measured with SWATH-MS and identified previously as exhibiting a significant strain-specific response after 6 and 12 weeks of high-fat diet *ad libitum* with a protein-protein interaction network from the String database. We used this information to perform a network enrichment analysis using the BioNet R package [14, 15] (see Methods). BioNet uses the topology of a protein-protein interaction network to calculate a minimum connected module that represents the maximum number of significantly changing proteins. We then performed pathway and go-term enrichment analysis in the obtained module to identify sub-modules of functional related proteins. With this we found several terms and pathways enriched in differentially abundant proteins, i.e. in proteins that reacted differently to a sustained high-fat diet depending on the genetic background [16, 17]. Gene ontology and pathway enrichment analysis was then performed on each of the functional modules (see Methods). Consistent with previous mRNA expression reports with similar experimental comparisons (Kirpich et al. [2011]), the identified functional modules were enriched in proteins involved in fatty acid metabolism, oxidation-reduction processes and stress-activated MAPK cascade, oxidation-reduction process, PPAR signaling pathway, biosynthesis of amino acids, fructose and mannose metabolism, cytochrome P450 metabolism, and peroxisome-related metabolic processes (**Figure 6** **and Supplementary Figure S5**).

**Figure 6.**
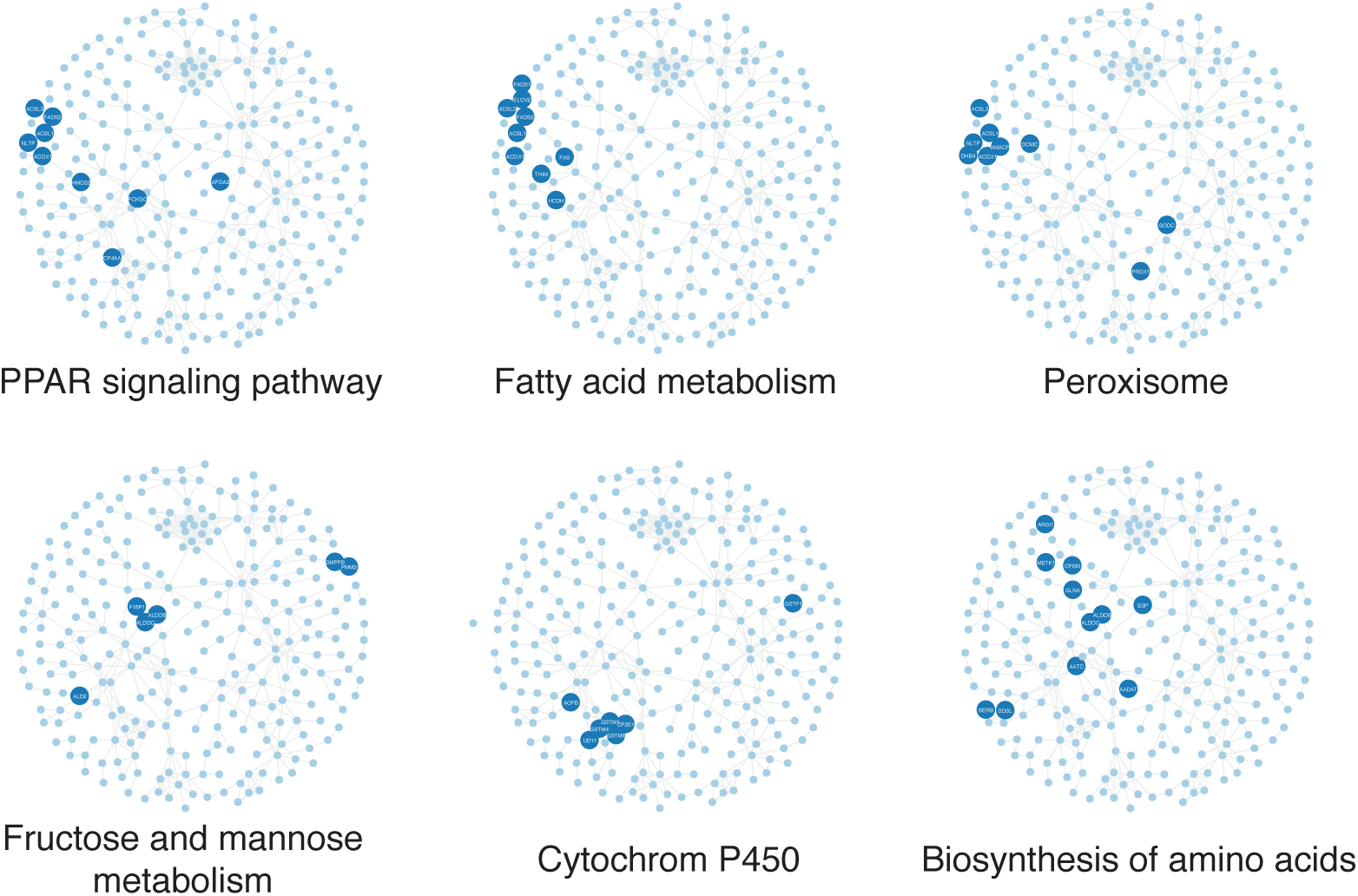
Network enrichment analysis. The adjusted p-values for the fold-change along time under high-fat diet for C57BL/6J (B6) and 129Sv (S9) mouse strains in the SWATH data are mapped to a protein protein interaction network, which is searched for functional modules enriched for highly varying nodes using BioNet (see Methods). The top scoring enriched sub-network is represented, with the GO gene sets and Kegg pathways found to be enriched in the functional module compared to the general protein protein interaction network as background (see Methods). Blue nodes correspond to protein belonging to the GO term or Kegg pathway and grey nodes are nodes belonging to the functional module. Node names are UniProt identifiers without the specie extension.

The analysis of functional modules was complemented by identifying proteins that showed (anti-)correlated quantitative patterns across the measured conditions. The goal of this step was to locate potential proteins subjected to mutual regulation with similar or opposed patterns. Taking as input the per-condition protein abundances estimated by MSstats, and the proteins present in the functional modules identified with BioNet, we calculated separately for each strain the Pearson correlation between all protein pairs connected in the functional network (**Supplementary Table ST3**). A ranked list based on the differences of the correlation coefficients between strains suggested pairs with a similar response to sustained high-fat diet in C57BL/6J mice but not in the 129Sv mouse strain, including the protein pairs CP4AA-UD11 involved in xenobiotic metabolism, AKC1H-3BHS5 related to steroid biosynthesis, and ITB1-ICAM1 with cell adhesion functions (**Figure 7**). Interestingly, we also found some functional protein pairs that exhibited a correlated response in 129Sv mice but not in the C57BL/6J mouse strain such as the protein pair AOFB-ADH1, which is also related to the xenobiotic metabolism.

**Figure 7.**
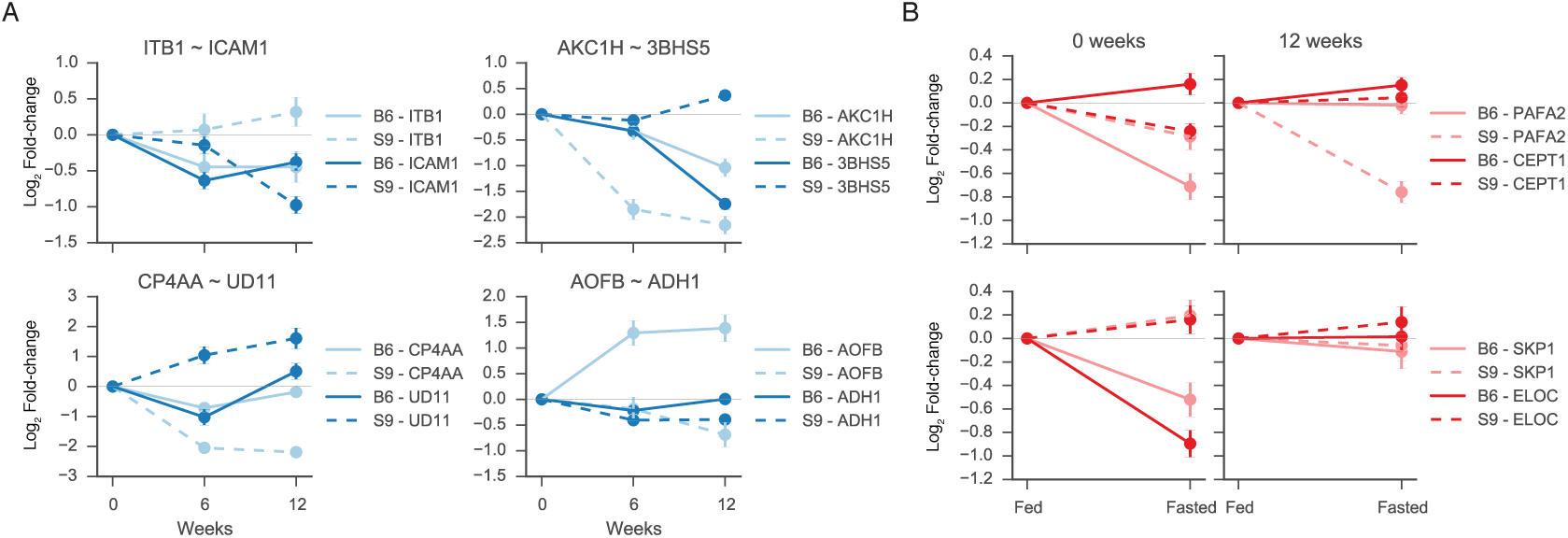
Protein fold-changes of functionally connected proteins that exhibit strain-specific responses (A) at different time points and (B) in the fed-to-fasted transition after a sustained high-fat diet. Protein relative abundance profiles of a subset of (anti)correlated protein pairs of functionally connected proteins as reported in the STRING functional interaction network, are shown for C57BL/6J (B6) and 129Sv (S9) mice in the SWATH-MS dataset.

### Influence of a sustained high-fat diet on the fed-fasting transition

Sustained high-fat diet is known to also alter the transient changes that normally occur when individuals shift from a fed to a fasted state. Therefore, apart from analyzing proteins with a significant strain-specific response upon a sustained high-fat diet, we were interested in elucidating how a persistent high-fat diet affected the fasting transition in C57BL/6J and 129Sv mice. To establish these effects we initially analyzed the changes occurring in the fed-to-fasted transition in the normal proteome, i.e. in the initial time point of 0 weeks, and compared the protein abundances of mice fed *ad libitum* and of mice fasted overnight between the two mouse strains. We then assessed whether the differences observed between strains when comparing fed-to-fasted states were altered when repeating the same fasting shift after 12 weeks of high-fat diet. In other words, we performed the two comparisons and selected those proteins that did not exhibit significant differences in the first comparison, but that exhibited a significant p-value the latter: i) (B6T0FAS–B6T0FED)–(S9T0FAS–S9T0FED), and ii) (B6T2FAS–B6T2FED)–(S9T2FAS–S9T2FED), with B6 representing in C57BL/6J mice, S9 representing 129Sv mice, and T0 and T2 corresponding to 0 and 12 weeks of high-fat diet. In this case *FED* referred to the treatment of mice were fed *ad libitum*, whereas *FAS* referred to mice fasted overnight. This analysis allowed us to identify 845 proteins showing a strain-specific abundance change during the fed-to-fasted transition after a sustained high-fat diet (**Supplementary Table ST4**). Protein changes occurring in strain-specific fed-to-fasted conditions were used to perform network enrichment analysis in a protein-protein interaction network, using the same approach as before (see Methods). The analysis of the interaction network resulted in the identification of functional modules related to fatty acid metabolism, to the ribosome and the proteasome, as well as proteins involved in valine, leucine and isoleucine degradation, in the PPAR signalling pathway, and propanoate metabolism (**Figure 7B**, **Supplementary Figure S6 and Supplementary Table ST5**).

## DISCUSSION

In systems biology, new knowledge is frequently generated by the measurement of molecular profiles in complex systems under differentially perturbed conditions where the data is used to support models and to generate or test new hypotheses. Most of these comparative studies have been carried out with transcriptomic experiments, but recent advances in reproducible, quantitative measurement of the proteins by mass spectrometry suggest that proteomics data could add complementary information much closer to the biological function.

In this work, we aimed to generate system-wide quantitative biological information related to the metabolic syndrome with the minimal required *a priori* expert knowledge for target protein selection. The metabolic syndrome was selected as a prototypical study case because of its medical relevance and complexity and concretely, we were interested in the links between diet and genetic background that trigger the onset of the metabolic syndrome. The metabolic syndrome is a combination of medical disorders that includes central obesity, insulin resistance, dyslipidemia, hypertension and hepatic steatosis, which predispose patients to several diseases such as diabetes, cardiovascular disease and cancer. Although extensive knowledge has accumulated about individual enzymes and regulatory proteins during the last years, the complex etiology of insulin resistance and its association to the accumulation of lipids in liver is not yet completely understood.

In this study we used two mouse strains, which exhibited different responses to a sustained high-fat diet under normal conditions, as a model to study the underlying mechanisms of the metabolic syndrome. Wild-type C57BL/6J mice showed altered physiological parameters such as higher body weight, increased fat accumulation in the liver, hyperglycemia and hyperinsulinemia, whereas 129Sv mice showed an increased resistance to these phenotypic alterations. We exposed these two phenotypically different mouse strains to long-term high-fat diet, and three different mass spectrometric acquisition approaches to generate a proteomics compendium consisting in three proteomics datasets: i) discovery shotgun, ii) SRM and iii) SWATH-MS.

We showed that targeted high-throughput approaches such as SWATH-MS are able not only to distinguish between biologically relevant factors in complex experimental designs, but also to reproduce the findings obtained with gold-standard methods in quantitative proteomics without the need of any previous expert knowledge for target protein selection. Indeed, using the targeted SWATH-MS dataset, we were able to qualitatively recapitulate previously reported biological findings [12] and, due to its high-throughput capabilities, we generated some additional insights that expanded beyond the proteins related to the central metabolism. reducing the risk of introducing protein selection bias, and opening the scope of investigation to not well-known systems.

## EXPERIMENTAL PROCEDURES

### Mice

Three-weeks old C57Bl/6J and 129Sv mice were obtained from Charles River Laboratories International, Inc. and maintained in a pathogen-free facility at the Institute of Molecular Systems Biology (ETH-Zurich), complying official ETH Zurich ethical guidelines. Mice from both strains were kept on a 12 h light/dark cycle and fed with a high-fat rodent diet (60 % fat content) for 0, 6 and 12 weeks according to the experimental design and as described previously [12].

### Sample preparation

PBS-perfused mouse livers were homogenized using RIPA-modified buffer (1% NP-40, 0.1% sodium deoxycholate, 150 mM NaCl, 1 mM EDTA, 50 mM Tris pH 7.5, protease inhibitors EDTA-free, 10 mM NaF, 10 mM sodium pyrophosphate, 5 mM 2-glycerophosphate) and a glass-glass tight douncer homogenizer (Wheaton Science Products, USA). Homogenates were centrifuged (20,000 g, 4 °C, 15 min) and the supernatant was collected and kept at 4 °C Pellets were resuspended with Urea-Tris buffer (50 mM Tris pH 8.1, 75 mM NaCl, 8 M urea, EDTA-free protease inhibitors, 10 mM NaF, 10 mM sodium pyrophosphate, 5 mM 2-glycerophosphate), and after sonication, they were centrifuged again (20,000 g, 4 °C, 15 min). The new supernatant was collected and mixed with the previous one whereas the resulting pellets were discarded. The total protein content was quantified with the BCA Protein Assay (Thermo Fisher Scientific, USA) using bovine serum albumin as standard. Aliquots of the homogenates were immediately prepared and stored at −80 °C Sample quality and concentration was double checked with SDS-PAGE and silver staining. 1 mg of total protein was mixed with 1 mg of the heavy-labeled reference proteome and the mix was precipitated overnight with 6 volumes of ice-cold acetone (16 h, 20 °C) for mass spectrometric analysis. The supernatant was discarded and pellets were dried and resuspended in freshly prepared digestion buffer (8 M urea, 0.1 M NH_4_HCO_3_). Samples were reduced with 12 mM dithiothreitol (30 min, 37 °C) and alkylated with 40 mM iodoacetamide (45 min, 25 °C) in the darkness. Samples were diluted with 0.1 M NH_4_HCO_3_ to a final concentration of 1.5 M urea and digested overnight at 37 *°C* with sequence grade trypsin (10 μg, Promega AG, Switzerland). After digestion, peptide mixtures were acidified to pH 2.8 with TFA and desalted with 500 mg Sep-Pak tC18 silica cartridges (Waters Inc., USA). Samples were dried under vacuum prior off-gel fractionation (24 wells, 310 pI strips) in a 3100 OFFGEL Fractionator (Agilent Technologies). Peptide mixtures collected in each well were pooled in six different fractions (A: wells 1- 2, B: wells 3-4, C: wells 5-8, D: wells 9-11, E: wells 12-18 and F: wells 19-24), acidified to pH 2.8 with trifluoroacetic acid and desalted with MacroSpin C18 silica columns (The Nest Group Inc., USA). Samples were dried under vacuum and re-solubilised to 1 μg/μl in 0.1% formic acid and 2% acetonitrile prior to MS analysis.

### Heavy-labeled reference proteome

A Hepa1-6 mouse cell line was obtained from the American Type Culture Collection (cat. CRL-1830), labeled with SILAC medium and used as a heavy-labeled reference proteome in all samples. Cells were grown at 37 °C and 5% CO2 in L-lysine-and L-arginine-depleted DMEM high glucose medium (Caisson Laboratories Inc) supplemented with 10% dialyzed FCS (BioConcept), 1% penicillin/streptomycin (Invitrogen) and 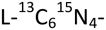 arginine and 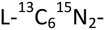 lysine (Sigma-Aldrich GmbH). Cells were cultured in 15-cm plates and passaged at 80% confluence until an incorporation of the heavy amino acids above 95% was achieved. Then, cells were washed with ice-cold PBS, scrapped from the plates and homogenised as described above.

### Shotgun MS measurements

Shotgun mass spectrometric measurements were performed on a LTQ-FT mass spectrometer (Thermo-Fischer Inc.) equipped with a nano-LC electrospray ionisation source. Fused-silica microcapillary columns (75 μm, New Objective Inc.) were packed with Magic C18 AQ 3 μm reverse-phase material (Michrom BioResources). Samples (2 μg) were analysed with a linear gradient from 5% to 30% of buffer B in 60 minutes at a flow rate of 300 nl/min using a Proxeon Easy-nLC 300 system (Thermo-Fisher Inc.). Buffer A: 98% H2O, 2% ACN and 0.1% formic acid; Buffer B: 2% H2O, 98% ACN and 0.1% formic acid. Three replicates for each time point were used in these measurements. MS2 data were analysed with the Sequest search engine (Sorcerer 4.0.4 build) against a mouse Uniprot database (v. dated 07/2009) containing all reviewed proteins and their corresponding decoy entries generated by reversing the amino acid sequences of the tryptic peptides. Search parameters were set to 50 ppm for the precursor mass tolerance, fully tryptic peptides, two miscleavages were allowed, and a false discovery rate of 1% based on PeptideProphet probability and decoy assignments [Keller et al., 2005]. Cysteine carbamidomethylation was set as a fixed modification, and methionine oxidation was used as a variable modification. Extracted MS1 peak intensities were used for protein quantification. Initially, mass spectrometric features were automatically detected, annotated, and aligned through the different samples using the software suite OpenMS (v1.7) [18]. Features completely missing in more than two thirds of the conditions were removed and with the rest a G-test with Williams correction was used to test for the pattern of missing values. Features not missing at random were imputed with the 10% of quantile values of peak intensities of each MS run. Finally, intensity values were transformed to a logarithmic scale and a constant normalization was performed in order to equalize the median peak intensities of reference heavy peptides between runs. Only normalized peak intensities corresponding to unique peptides were retained for quantitative analysis. Protein-level quantification and testing for differential abundance were performed using a linear mixed-effects model (from the R package MSstats v2.3.7) with terms for the conditions, biological replicates, peptides, MS runs and corresponding interactions [19, 20], and reduced scope of biological replication A list of p-values was calculated for each comparison, and was adjusted to control the False Discovery Rate (FDR) at 1%. Original scripts are available in https://github.com/saezlab/liverx.

### SRM Measurements

SRM measurements were performed as previously described [Sabidó et al. 2013]. Briefly, for each selected protein, unique tryptic peptides were selected for SRM quantification based on previously observed peptides in shotgun proteomics experiment as reported in publicly accessible proteomic data repository PeptideAtlas [21] or based on bioinformatic prediction when no peptides had been observed. The selected peptides were synthesized and analyzed via SRM-triggered MS2 acquisition on the triple quadrupole mass spectrometer to obtain the corresponding full fragment ion spectra. The most intense fragment ions from the full fragment ion spectra were used for SRM MS measurements on a hybrid triple quadrupole-ion trap mass spectrometer (4000QTrap; ABI/MDS-Sciex) equipped with a nanoelectrospray ion source. Chromatographic separations of peptides were performed a 15-cm analytical column (75 μm) packed with a Magic C18 AQ 5 μm resin (Michrom BioResources). A linear gradient from 5 to 35 % of buffer B in 35 min at a flow rate of 300 nl/min was used (Tempo nanoLC system; Applied Biosystems, USA). Buffer A: 98% H2O, 2% ACN and 0.1% formic acid; Buffer B: 2% H2O, 98% ACN. Acquired SRM traces were evaluated based on co-elution, shape similarity and intensity rank between the endogenous and isotopically-labeled reference peptide digest, and the reference MS/MS spectra. Peptide areas were normalized based on the isotopically-labeled reference peptide areas, and protein abundance changes were inferred using a linear mixed-effects model for protein quantification as implemented in MSstats v2.3.7 using reduced scope of biological replication [19]. A list of p-values was calculated for each comparison, and was adjusted to control the False Discovery Rate (FDR) at 1%. Original scripts are available in https://github.com/saezlab/liverx.

### SWATH-MS Measurements

SWATH MS was performed on a 5600 TripleTOF mass spectrometer as previously described [8]. The mass spectrometer was tuned to allow a quadrupole resolution of 25 Da/mass selection, and a set of 32 overlapping windows was constructed covering the precursor mass range of 400-1200 Da. The effective isolation windows can be considered as being 399.5-424.5, 424.5-449.5, etc. plus the potential overlap left from the nominal window transmission. SWATH data extraction were performed using OpenSWATH [22]. Extracted fragment ion areas were used to infer protein abundance changes using a linear mixed-effects model for protein quantification as implemented in MSstats v2.3.7 using reduced scope of biological replication [19]. A list of p-values was calculated for each comparison, and was adjusted to control the False Discovery Rate (FDR) at 1%. Original scripts are available in https://github.com/saezlab/liverx.

All mass spectrometry proteomics data have been deposited to PRIDE (in process).

### Statistical analysis

Basic statistical analyses and plotting were done in Python (v2.7.6). For the analysis of correlation patterns, Pearson correlation coefficients between protein estimates in pairs of conditions were computed using proteins with an FDR-adjusted p-value below 0.05 in at least one condition. The principal components analysis was performed using the *PCA* Scikit-learn (v0.16.1) function. All details are described in the original scripts available in https://github.com/saezlab/liverx.

### Network analysis

We used the String v10 [16, 17] mouse interaction network downloaded on the 18^th^ of June 2015 and filtered to exclude interactions setting String’s stringency score to 800 (out of maximum score of 999) [16]. String assembles protein-protein interactions from multiple resources, including genomic context, high-throughput experiments, or from previous knowledge databases, e.g. PubMed, and ranks them according to their evidence. Only interactions between nodes with a mapped protein measurement were used. No self-loops were allowed.

The network enrichment analysis was done using the *BioNet* R package and *Heinz* algorithm (R v3.1.2, BioNet v1.26.1, Heinz v2.0) [14, 15] using a 5% FDR threshold. BioNet allows the identification of a minimum connected protein-protein module that maximises the number of proteins changing significantly in abundance using the Heinz algorithm [14, 15].

Network enrichment was performed for the two biological questions of interest, i.e. time effects of a sustained high-fat diet on different genetic backgrounds and the Influence of a sustained high-fat diet on the fed-fasting transition. The network for high-fat diet on different genetic backgrounds contains 273 nodes and 530 edges and for the fed-fasting transition contains 442 nodes and 1323 interactions. Each sub-network was independently used for gene set enrichment analysis using a hypergeometric test (with the full network as background) using *hypergeom* function from Scipy module (v0.15.1). Kegg pathways were obtained using Bioservices (v1.3.6) [23] Kegg service, Kegg gene identifiers were converted to UniProt ids using Kegg BioService internal converter function *conv*. GO terms were downloaded on the 15^th^ of June 2015 from UnipProt-GOA. Pearson correlations between protein pairs were calculated separately for each mouse strain by concatenating the fed and fasted profiles. Original scripts are available in https://github.com/saezlab/liverx.

## SUPPLEMENTARY INFORMATION

**Supplementary Figure S1**

A) Overlap of measured proteins by SWATH, SRM, and shotgun proteomics. B) Density curves and C) violin plots representing the log2-fold-change distributions for the different mass spectrometric datasets acquired. B) and C) overlay values for the following tested comparison between conditions: i) B6T1FED–B6T0FED, ii) B6T2FED–B6T0FED, iii) B6T2FED–B6T1FED, iv) S9T1FED–S9T0FED, v) S9T2FED–S9T0FED, and vi) S9T2FED–S9T1FED). B6 represents C57BL/6J mice, and S9 represents 129Sv mice. T0, T1 and T2 correspond to 0, 6 and 12 weeks of high-fat diet. FED stands for mice that were fed *ad libitum* (treatment).

**Supplementary Figure S2**.

A) Volcano plots representing the-log10 of the adjusted p-value versus the log2-fold-change protein estimates for the overlapping proteins in discovery shotgun, SWATH-MS and SRM datasets. B) Density curves and C) violin plots representing the log2-fold-change distributions for the different mass spectrometric datasets acquired for the overlapping proteins.

**Supplementary Figure S3**

Pearson correlation of log-fold changes of proteins between SRM, SWATH, and shotgun technologies for the subsets of proteins quantified with two acquisitions.

**Supplementary Figure S4**

Fold-changes (log2-scale) corresponding to proteins involved in the main metabolism after 0, 6 and 12 weeks of sustained high-fat diet for C57BL/6J (B6) and 129Sv (S9) mice in the SWATH-MS quantitative dataset for both fed and fasted condition.

**Supplementary Figure S5**

Gene ontology and pathway enrichment analysis of the protein network obtained from proteins that with a significant strain-specific response in at least one time point after 0, 6 and 12 weeks of sustained high-fat diet in the SWATH-MS quantitative dataset.

**Supplementary Figure S6**

Gene ontology and pathway enrichment analysis of the protein network obtained from proteins that in the SWATH-MS quantitative dataset exhibit strain-specific changes in the fed-fasted transition after a sustained high-fat diet.

**Supplementary Table ST1**

Extracted raw areas for precursor and fragment ions quantified in the discovery shotgun, SRM and SWATH-MS datasets.

**Supplementary Table ST2**

Statistical analysis of SWATH-quantified proteins to detect proteins that change in abundance after 0, 6 and 12 weeks of high-fat diet in for C57BL/6J (B6) and 129Sv (S9) mice.

**Supplementary Table ST3**

Correlation of estimated abundances between proteins changing significantly over time among mouse strains and that are linked according to the STRING functional network.

**Supplementary Table ST4**

Statistical analysis of SWATH-quantified proteins to the effects of a persistent high-fat diet in the fasting transition in C57BL/6J and 129Sv mice.

**Supplementary Table ST5**

Correlation of estimated abundances between proteins changing significantly in the fed fasted transition among mouse strains and that are linked according to the STRING functional network.

## AUTHOR CONTRIBUTIONS

RA conceived the project; ES designed the experiments; ES, YW, LB and TP performed the experiments; CT, EG and JSR did the computational analyses; TC, VC, OV contributed to the data processing; CT, EG, ES, YW, JSR and RA interpreted the data; CT, ES, EG, RA and JSR wrote the manuscript.

## ACKNOWLEDGEMENTS

Support for this study was provided by the European Union via the ERC (European Research Council) advanced grant ‘Proteomics v3.0’ (grant# 233226) to RA, and the LiverX program of the Swiss Initiative for Systems Biology (SystemsX) to ES, YW and RA.

